# Survey of *Peristenus* spp. (Hymenoptera: Braconidae) parasitoids in the Monterey Bay region of California

**DOI:** 10.1101/2025.11.26.690901

**Authors:** Tucker Morrison, Diego J. Nieto, Jason Buhler

## Abstract

Two European parasitoids, *Peristenus relictus* and *Peristenus digoneutis*, were introduced into California to improve the management of *Lygus hesperus*. While *P. relictus* quickly established, *P. digoneutis* has only rarely been recovered. After a renewed release effort, a field survey was conducted in an attempt to recover *P. digoneutis*. Collections of *Lygus hesperus* nymphs were made from strawberry production areas and associated alfalfa trap crops in the Monterey Bay region in 2023. Parasitoid larvae recovered from dissected nymphs were identified using species-specific molecular markers. All successfully assayed specimens were confirmed as *P. relictus*. Furthermore, all reared parasitoids from similarly collected *Lygus* nymphs were identified as *P. relictus*. This survey corroborates previous findings indicating that *P. relictus* is the primary nymphal parasitoid in California and fails to provide evidence that *P. digoneutis* has established on the central coast.

## Introduction

*Lygus hesperus* (Hemiptera: Miridae) is a key pest of strawberry and other specialty crops in California. In response, two European braconid parasitoids, *Peristenus relictus* (Ruthe) and *Peristenus digoneutis* (Loan), were introduced into California in 1998 (Pickett et al., 2007). These parasitoids were initially released in the Sacramento Valley and later released in the Monterey Bay region from 2002-2006 (Pickett et al., 2007, 2009). *Peristenus relictus* proceeded to disperse widely from release sites and is now established in all three primary strawberry growing regions in coastal California (Pickett et al., 2013; Nieto et al., 2020).

In contrast, *P. digoneutis* has not been recovered in Sacramento after its initial colonization (Pickett et al., 2007). On the central coast, overwintered collections of *P. digoneutis* were similarly absent immediately following releases from 2002-2006. Subsequent surveys within the Central Valley (Pickett et al., 2013) and along the central coast (Nieto et al., 2020) also lacked *P. digoneutis*. As a result, a renewed effort to source a population of *P. digoneutis* from a region in France with a similar climate to coastal California led to a second round of releases in California during the 2010’s. Here we provide a survey of *Lygus* nymphal parasitoids conducted in the Monterey Bay region in 2023 that includes *Peristenus* species identifications using both rearing and PCR methods.

## Methods

### *Lygus* nymph collection

*Lygus* nymphs were collected in and around strawberry fields and associated alfalfa trap crops in Santa Cruz and Monterey Counties in the summer of 2023. In several cases, collections were taken from sites where *P. digoneutis* were previously released. Nymphs were collected using either a standard sweep net or a hand-held bug vacuum (Stihl SH 56 C-E). Each sample consisted of between 50 and 200 successive sweeps or suctions. A single such collection of *Lygus* nymphs was taken per site to rear out either *Lygus* or *Peristenus* sp. adults (Table 1). For molecular analyses, *Lygus* nymphs were provided by either three-to-four samples taken per location (Table 2) or were gleaned from broader (i.e., unrelated) collection efforts (Table 3).

**Table 1:** *Peristenus* sp. reared from *Lygus* nymphs collected in the Monterey Bay region in 2023.

| date | sample location | plant | # sweeps | species (# reared) |
| --- | --- | --- | --- | --- |
| 19 June | Salinas-1 | alfalfa | 100 | 0 |
| 14 July | Watsonville-1 | weeds | 50 | *P. relictus* (53) |
| 15 July | Salinas-2 | sweet alyssum | 50 | 0 |
| 7 August | Salinas-1 | alfalfa | 200 | *P. relictus* (24) |
| 7 August | Watsonville-2 | sweet alyssum | 200 | *P. relictus* (45) |
| 10 August | Salinas-3 | alfalfa | 200 | *P. relictus* (23) |
| 16 September | Salinas-3 | alfalfa | 200 | *P. relictus* (4) |
| 18 September | Salinas-4 | alfalfa | 200 | *P. relictus* (4) |
| 21 September | Salinas-3 | alfalfa | 200 | *P. relictus* (26) |
| 22 September | Salinas-2 | sweet alyssum | 200 | *P. relictus* (7) |
| 25 September | Watsonville-2 | sweet alyssum | 200 | *P. relictus* (13) |
| 25 September | Salinas-2 | sweet alyssum | 200 | *P. relictus* (33) |

**Table 2:**
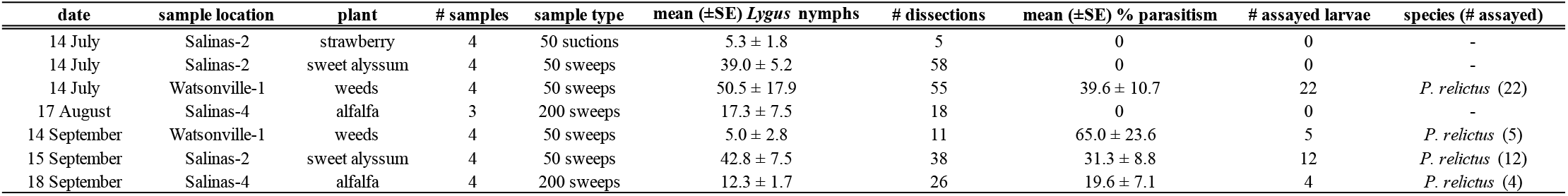
Parasitism rates and PCR-based identification of *Peristenus* larvae from *Lygus* nymphs collected in the Monterey Bay region in 2023.

**Table 3:** PCR-based identification of *Peristenus* larvae harvested from *Lygus* nymphs collected in the Monterey Bay region in 2023.

| month | sample location | plant | sample type | # sample dates | # assayed larvae | species (# assayed) |
| --- | --- | --- | --- | --- | --- | --- |
| July | Salinas-3 | alfalfa | 200 sweeps | 2 | 30 | *P. relictus* (30) |
| August | Watsonville-3 | alfalfa | 200 sweeps | 1 | 1 | *P. relictus* (1) |
| August | Salinas-3 | alfalfa | 200 sweeps | 3 | 32 | *P. relictus* (32) |
| August | Salinas-3 | strawberry | 100 suction | 2 | 10 | *P. relictus* (10) |
| August | Salinas-4 | weeds | 200 sweeps | 1 | 1 | *P. relictus* (1) |
| August | Watsonville-2 | sweet alyssum | 100 suction | 1 | 9 | *P. relictus* (9) |
| August | Salinas-5 | alfalfa | 200 sweeps | 2 | 41 | *P. relictus* (36) |
| August | Salinas-1 | alfalfa | 200 sweeps | 4 | 36 | *P. relictus* (25) |
| August | Salinas-1 | strawberry | 100 suction | 3 | 3 | *P. relictus* (3) |
| September | Salinas-3 | alfalfa | 200 sweeps | 4 | 30 | *P. relictus* (27) |
| September | Salinas-5 | alfalfa | 200 sweeps | 2 | 2 | *P. relictus* (2) |
| September | Salinas-1 | alfalfa | 200 sweeps | 3 | 13 | *P. relictus* (13) |

Collected nymphs were transported to the Driscoll’s Inc. Entomology laboratory, where they were further processed. Reared adult parasitoids were identified using Goulet and Mason (2006).

### Larval extraction and molecular identification

Nymphs were counted and dissected under a stereomicroscope within 24 hours of collection. Third-, fourth-, and fifth-instar nymphs were selected for dissection, as these stages are more likely to contain parasitoid larvae. Dissection methods used here are described by Nieto et al. (2025). Dissected parasitoid larvae were stored individually in 0.2 mL PCR tubes containing 95% ethanol at -20°C until DNA extraction.

Total DNA was extracted from individual parasitoid larvae using the DNeasy Blood and Tissue Kit (Qiagen) following the manufacturer’s protocol with all reagent volumes reduced to one-third to accommodate the specimen’s small size. Final elutions used 50 μL of molecular-grade water. Species identifications used species-specific PCR primers developed by Gariepy et al. (2005) targeting the ITS1 region. The primer set consists of species-specific reverse primers (digF1096 for *P. digoneutis*, styF1230 for *P. relictus*) and a conserved forward primer (Per R1), amplifying diagnostic fragments of 515 bp for *P. digoneutis* and 330 bp for *P. relictus*.

To perform PCR, reaction mixtures (25 μL) contained 12.5 μL OneTaq 2X Master Mix (New England Biolabs), 0.5 μM each primer, 2 μL template DNA, and molecular-grade water. Each DNA sample was amplified in 2 separate reactions using each primer pair.

To amplify DNA, the initial denaturation was set at 94°C for 120 s, followed by 35 cycles of 94°C for 45 s, 54°C for 45 s, and 72°C for 60 s, with a final extension set for 5 min at 72°C. PCR products were visualized on 1.5% agarose gels in TBE buffer, stained with GelRed, and visualized under UV illumination. Product identity was confirmed by comparison to positive controls (DNA from morphologically identified adult *P. relictus* and *P. digoneutis*) and a 100 bp DNA ladder.

### Results and discussion

Of the 403 *Lygus* nymphs that were successfully reared in 2023, 171 reached adulthood. The remaining 232 nymphs produced eclosed adult parasitoids, all of which were identified as *P. relictus* (Table 1).

A total of 251 *Peristenus* sp. larvae were assayed across all collection sites in 2023 (Tables 2-3). Of these, 232 were successfully assayed; 19 larvae failed to amplify and were excluded from analysis. All successfully assayed larvae were identified as *P. relictus* based on the presence of the 330 bp amplicon in reactions using the *P. relictus*-specific primer pair. No specimens produced the 515 bp amplicon diagnostic of *P. digoneutis* in reactions using the *P. digoneutis*-specific primer pair. These results were consistent across all sampled locations and collection dates.

*Peristenus relictus* was collected from all sampled coastal locations, confirming its widespread distribution in this strawberry-growing region. Conversely, the absence of *P. digoneutis* recoveries indicates that the more recent releases of this parasitoid may not have led to a successful colonization on the central coast. If so, incompatible climatic conditions relative to overwintering needs may be the cause.

